# 3D Transcranial ultrasound localization microscopy reveals major arteries in the sheep brain

**DOI:** 10.1101/2024.02.28.582489

**Authors:** Antoine Coudert, Louise Denis, Arthur Chavignon, Sylvain Bodard, Mikael Naveau, Palma Pro Sistiaga, Romaric Saulnier, Cyrille Orset, Denis Vivien, Christine Chappard, Olivier Couture

## Abstract

**Objective:** Stroke, a leading cause of mortality and disability globally, demands swift and accurate diagnosis for effective treatment. Although MRI and CT scans serve as conventional methods, their accessibility remains a challenge, prompting exploration into alternative, portable, and non-ionizing imaging solutions like ultrasound with reduced costs. While Ultrasound Localization Microscopy (ULM) displays potential in high-resolution vessel imaging, its 2D constraints limit its emergency utility.

**Materials and Methods:** This study delves into the feasibility of 3D ULM with multiplexed probe for transcranial vessel imaging in sheep brains, emulating human skull characteristics. Three sheep underwent 3D ULM imaging, compared with angiographic MRI, while skull characterization was conducted in vivo using ultrashort bone MRI sequences and ex vivo via micro CT.

**Results and conclusions:** The study showcased 3D ULM’s ability to highlight vessels, down to the Circle of Willis, yet within a confined 3D field-of-view. Future enhancements in signal, aberration correction, and human trials hold promise for a portable, volumetric, transcranial ultrasound angiography system.

**Summary statement:** 3D Ultrasound localization microscopy, using a low-frequency matrix probe, enables transcranial reconstruction of the main vessels in sheep brains.

## Introduction

Cerebral circulation supplies nutrients and oxygen to more than 120 billion neurons^1^. However, during a stroke, blood flow is reduced and 1.9 million neurons are lost per minute on average^2^. As it occurs in 12 million patients per year, this makes stroke one of the major causes of mortality and the leading cause of acquired disability^3^. Rapid access to treatment reduces drastically morbidity and mortality^4^, but the therapeutic strategy depends strongly on the nature of the stroke. In 87% of the cases, strokes are ischemic, i.e. due to a blood clot, and they require thrombolysis or thrombectomy^5^. In the remaining cases, strokes are mainly due to a vessel rupture, leading to a hemorrhage that require bleeding control.

As the symptoms are often indistinguishable between ischemic and hemorrhagic stroke, medical imaging is used for their diagnosis. MRI and CT are preferred due to their various contrast mechanisms sensitive to blood flow, perfusion, or tissue alteration^6^. However, swift access to these imaging modalities remains limited. To reduce the time interval before treatment, mobile stroke units equipped with CT scanners have been evaluated in the USA and Germany and shown to reduce treatment intervention time significantly^7,8^. Nevertheless, such an approach is restricted by the weight, cost, and ionizing radiation of the CT scanner.

Ultrasound has the advantage of being an easily transportable, relatively inexpensive and non-irradiating technology with few contraindications. It could therefore be an interesting candidate for ambulance-based diagnosis. However, ultrasound of the brain is severely hampered by the presence of the skull. The three skull bone layers, i.e. inner/outer cortices with diploe between them, cause aberration and attenuation of sound waves^9–12^. As the attenuation is worsened at higher frequencies^13^, transcranial Doppler is currently performed around 2 MHz, which limits the resolution of ultrasound imaging and its reliability as a diagnostic device^14^.

In recent years, super-resolution ultrasound imaging has been developed^15–17^. In particular, Ultrasound Localization Microscopy (ULM) allows the observation of vessels at a resolution below the diffraction limit. Thanks to an intravenous microbubble’s injection used in clinical routine, we can observe these microbubbles progression through the vasculature with ultrasound imaging. Given the separation of each microbubble, their location can be established within a few microns. ULM maps can then be reconstructed by cumulating millions of individual microbubble tracks, yielding to vessels density and velocity information.

ULM has been applied in a preclinical context on many organs, such as the spinal cord, kidneys, liver tumors, and the ovaries^18–22^. In our group, we have shown the ability of ULM to map kidney’s microcirculation in humans with a clinical US scanner^23–25^. The brain^16^ has also been studied in rodents for Alzheimer’s disease^26^ and on functional imaging^27^ or stroke^28^. In case of human brain, it has shown its ability to overcome the penetration/resolution trade-off in the human brain and to image the vessels, including an aneurysm^29^.

In previous study, ULM in humans has been restricted to 2D imaging. However, such dimensional constraint imposes severe limitations to the generalization of this technique and portable ultrasound in general. Firstly, appropriate quantification of out-of-plane flow or motion is not possible. Secondly, only a limited number of microbubble events can be intercepted in a given period and restriction to 2D imaging ultimately impairs resolution^30^. Thirdly, the selection of the imaging plane requires a high level of radiological expertise at the bedside which is not compatible with the emergency situation^31,32^. This is particularly problematic for ultrasound imaging through the temporal window. This acoustic window is mainly used in clinics due to its lower attenuation because of its thinner thickness, i.e. between 1.5 and 6mm, and its low porosity^33^. But even if this temporal window is highly appropriate in the context of strokes, i.e. 51% of ischemic strokes occur around the middle cerebral artery visible through the temporal window^34^, probe positioning remains highly user-dependent due to the small acoustic window area.

To avoid these issues and in particular plane-selection with portable scanners, ULM should be performed in 3D. Several methods exist. The first possibility is to use an array probe and move it to image several slices of the same volume^16,35^. However, this method is unsuitable for microbubble tracking due to its low volume rate. The second is the use of a matrix probe, which has elements positioned both in the lateral and elevation directions, i.e. 32 × 32 for a 1024 elements probe. In the case of a 1024 elements fully-addressed independent electronic emission and reception channels, the device transportability is affected by the complexity of the electronic setup^36–39^. One option to avoid the fully addressed probe is to keep the same aperture, but reduce the number of elements by using a sparse array^40^. However, the reduction in signal-to-noise ratio is incompatible with transcranial imaging. On the other hand, row-column matrix can reduce the number of channels from N^2 to 2N+1 channels^41,42^, but beamforming and acquisition time needs to be adapted for such geometry. Another option is to use a multiplexer which allows successive allocation of numerous elements on a single channel. Multiplexing reduces volume rate but provides equivalent imaging to the fully assigned probe^43^. Such approach has been used in small animal models of stroke^28^ to perform 3D ULM with a high-frequency probe. Today, it remains to show the ability of this 3D ULM imaging method to image blood vessels in the case of a larger brain and a thicker skull which would be closer to the clinical context in humans.

In this study, we proposed to image sheep’s brain with the 3D ULM multiplexed method. This model was chosen because the thickness of the sheep’s skull, around 5 mm, is similar to that of the temporal region of the human skull^33,44,45^. 3D imaging was carried out with a low-frequency matrix probe of 1.5 MHz to reduce the attenuation of the skull. The loss of resolution was then compensated by ULM. 3D ULM was compared with angiographic MRI (MRA) and angiographic-CT. Finally, a characterization of the skulls was carried out in acoustic terms from Zero Time Echo (ZTE) MRI and in terms of microarchitecture from micro-computed tomography (micro-CT) imaging.

## Materials and Methods

### A. Experimental procedure

#### a. Animal procedure

This study was carried out in accordance with the ARRIVE guidelines, the European Directives and the French Legislation on Animal Experimentation (protocol authorization APAFIS #32738), and also approved by the local ethic committee (CENOMEXA, N°54). A total of 6 domestic sheep (Ovis aries weighting 35-40 kg, aging 15-16 months) were included, comprising the first 3 sheep for the establishment of the experimental protocol. Results presented here are from the 3 remaining sheep 4, 5 and 6 noted M4, M5 and M6. During the surgical and imaging procedures, general anesthesia was maintained.

Female sheep were sedated with ketamine (12 mg/kg) and xylazine (1 mg/kg) to allow placement of a venous line. Propofol (100 µg/kg) was then administered and endotracheal intubation was performed. During the imaging procedures, anesthesia was maintained by administration of sevoflurane (1.5-2.0 %) in air. The respiratory rate and tidal volume were adjusted to maintain physiological limits. Cardiac activity and oximetry were controlled, arterial pressure was maintained between 80 and 120mm Hg. Each animal’s body temperature was kept at 38°C with an air-heating blanket.

All procedures lasted 5 hours from sedation to final wake up in the original box stabling. After 3 imaging sessions conducted over one or two weeks and including ultrasound and MRI sequences, animals were euthanized with barbiturate overdose (pentobarbital 0.2 g/kg) after 2 minutes of 8% sevoflurane inhalation. Death was assessed by < 20mmHg arterial pressure and no cardiac activity.

After death, the part of the skull below the probe was removed, cleaned and preserved in physiological saline with a drop of bleach. It was then fixed in a bath of formalin-free fixative F13 (Morphisto, Germany) for one day then dried and placed under vacuum for micro-CT imaging.

#### b. CT-scan acquisitions

CT-scan was performed with VCT Discovery (GE, USA) on the sheep M6 only. A first sequence was done to highlight the skull (Helical slice thickness 0.625mm, 120 kV, 600 mA.S, Kernel BONEPLUS GE reconstruction) and the second one was performed with an iodine injection (iomeprol 400 mg/ml, injection 30 ml, 4 ml/s +20 ml of saline solution) for angiography (Helical slice thickness 0.625 mm, 120 kV, 600 mA.S, spiral pitch ratio 0.5, Kernel SOFT GE reconstruction).

#### c. MRI acquisitions

T2 MRI, and angiographic MRI (MRA) sequences were done on all the sheep using a 3T MRI Scanner (GE SIGNA PREMIER) in a thermo-regulated room, i.e. around 20°C (Axial Propeller, TR=5539ms, TE=165.744ms, slice thickness 2mm, 31 slices, in-place resolution 0.375×0.375). A ZTE MRI sequence was used to measure the bone thickness (3D coronal radial acquisition, FOV=16.0×16.0 cm², Matrix=512×512, Slice thickness = 1 mm, TR=600 ms/TE=0.016 ms). Finally, following a gadolinium (DOTAREM or CLARISCAN Gé, 0,5 mmol/mL solution for injection, gadoteric acid) injection through the femoral vein (0.2 ml/Kg 4 ml/s injection + 20 ml saline solution, 3 ml/s) a 3D arterial angiography was acquired (3D axial fast gradient echo, FOV=18.5×18.5 cm², Matrix=512×512, Slice thickness=0.6 mm, TR=4.74 ms/TE=1.696 ms, phase 1: passage through the carotid arteries and phase 2: 60 sec after phase 1).

#### d. Ultrasound acquisitions

The probe was placed above the shaved head of the sheep, after locating the thinnest area on the ZTE MRI sequence. To ensure appropriate placement over the brain, the probe was placed above the crest of the frontal bone pressed against the parietal bone, thanks to a contrast Power Doppler sequence, i.e. with 1 mL injection of ultrasound contrast agent (Sonovue, Bracco, Italy, sulfur hexafluoride, 8 µl/ml). High-volume rate imaging was performed using a research ultrasound scanner Vantage 256 (Verasonics, Kirkland, USA) and a matrix probe centered at 1.5 MHz (Vermon, Tours, France). The probe was composed of 1024 elements of 500 µm pitch, aligned on a 32×35 grid. The 9th, 18th, and 27th lines were inactive and 300 µm wide, creating 4 independent panels. The 1024 elements were connected to the 256 channels of the ultrasound scanner using a 4-to-1 multiplexer (Verasonics UTA 1024-MUX Adapter), which had 256 independent switches. Namely, the switch “i” was successively linked with elements “i”, “i+256”, “i+512”, and “i+768”. A switch could change its position, allowing emission with one element and receiving with another from this list.

A hybrid sequence of 3 cylindrical waves emitted by all the probe elements at the same time and 2 spherical waves decomposed by panel have been used to reach an imaging volume-rate of 209 Hz. Each pulse was composed of 2 cycles and the voltage was adapted to each sheep (Table 1). The injection of microbubbles (Sonovue, Bracco, Italy, sulfur hexafluoride, 8 µl/ml) for ULM depended on the animal. This consisted of injecting a 1.5 to 2 ml/minute for the all ULM acquisition duration, i.e. from 2.5 minutes to 6.5 minutes.

**Table 1.** Parameter of ULM acquisitions.

|  | Concentration | Number of injections | Number of frames | Distance virtual sources to probe (Wavelength) | Virtual sources (angle °) | Voltage |
| --- | --- | --- | --- | --- | --- | --- |
| M4 | 2 ml/min | 6 | 28000 | 60 | 5° | 60V |
| M5 | 1.5 ml/min | 6 | 80000 | 50 | 5° | 50V |
| M6 | 1.5 ml/min | 10 | 80000 | 30 | 7.5° | 60V |

#### e. Micro-CT acquisitions

After sheep euthanasia and extraction of the skull samples, samples were scanned with the Skyscan 1176 (Bruker, Kontich, Belgium) at 18 µm of resolution with isotropic voxel size. The micro-CT images were acquired with the following parameters: 90 kV X-ray tube voltage, 278 µA X-ray current, Filter: 0.3 mm Cu+0.5 mm Al, 420 ms integration time, frame averaging n=5, and 0.3° angular rotation step through 180° of rotation. The 2 cylindrical calibration phantoms provided by the manufacturer 3 cm in diameter were acquired at the same time for each specimen, they are composed of known concentrations of hydroxyapatite (HA) in epoxy resin of 0.250 gHA/cm^3^ and 0.750 gHA/cm^3^ (QRM, Freiburg, Germany). Three to five scans were required for specimens; they were reconstructed by a modified Feldkamp algorithm and joined by the post-processing alignment software NRecon (version 1.7.1.0).

### B. Data treatments

#### a. ULM Treatments

High volume rate ultrasound images were reconstructed with a delay-and-sum beamforming algorithm into a 300 × 300 × 600 mm^3^ volume with 0.5 × 0.5 × 0.5 mm^3^ voxels. Tissue signal was removed by the subtraction of the signal from a moving average (2 frames). After detection with a local maxima algorithm, microbubbles were localized by three-dimensional radial symmetry algorithm using an 11 pixels wide PSF (diameter 5.5 mm)^39^. Microbubbles with an amplitude of 10.5 dB above the surrounding noise floor were kept and tracked by the Hungarian method^46^ (<u>simpletracker - File Exchange - MATLAB Central</u> (mathworks.com)) with a minimal persistence time between 5 and 7 frames and a maximum pairing distance of 1.5 mm between two successive microbubble’s positions, i.e. maximum velocity of 350 mm/s (Fig. 2.A). All the tracks were then binned on a 0.1 mm wide voxel matrix to obtain an ULM density map, i.e. microbubbles count per voxel (Fig. 2.B, C)

**Figure 1.**
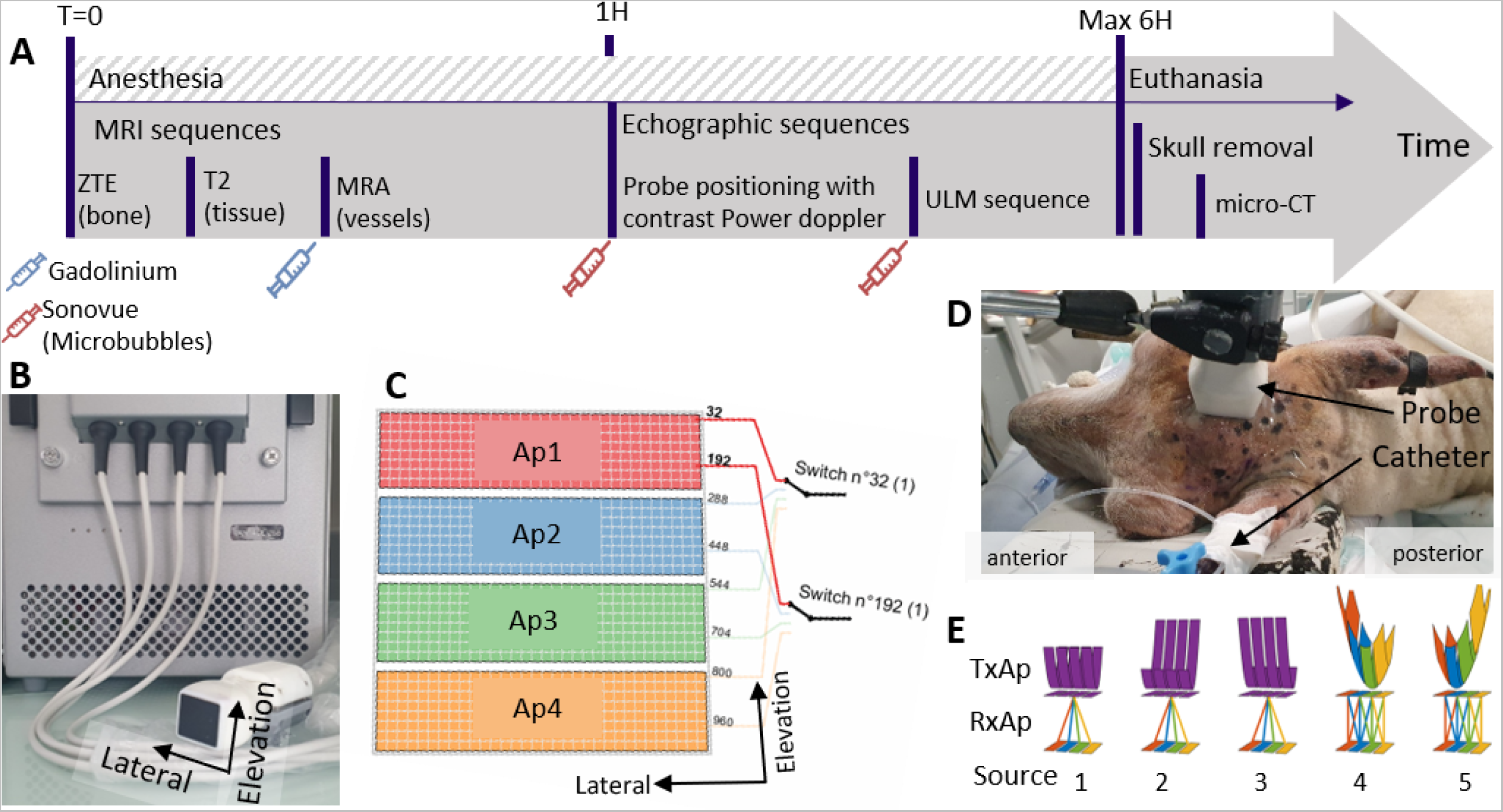
Experimental procedure. (A) Chronology of experimentations. (B) Matrix probe with the ultrasound scanner Verasonics Vantage 256. (C) Schematic of the grid of the 1024 elements of the probe which formed 4 apertures. (D) Probe positioning on the sheep’s head. (E) Schematic of the sequence. 3 full cylinder emissions and 2 spherical waves decomposed in four parts that’s leads to 32 emissions.

**Figure 2.**
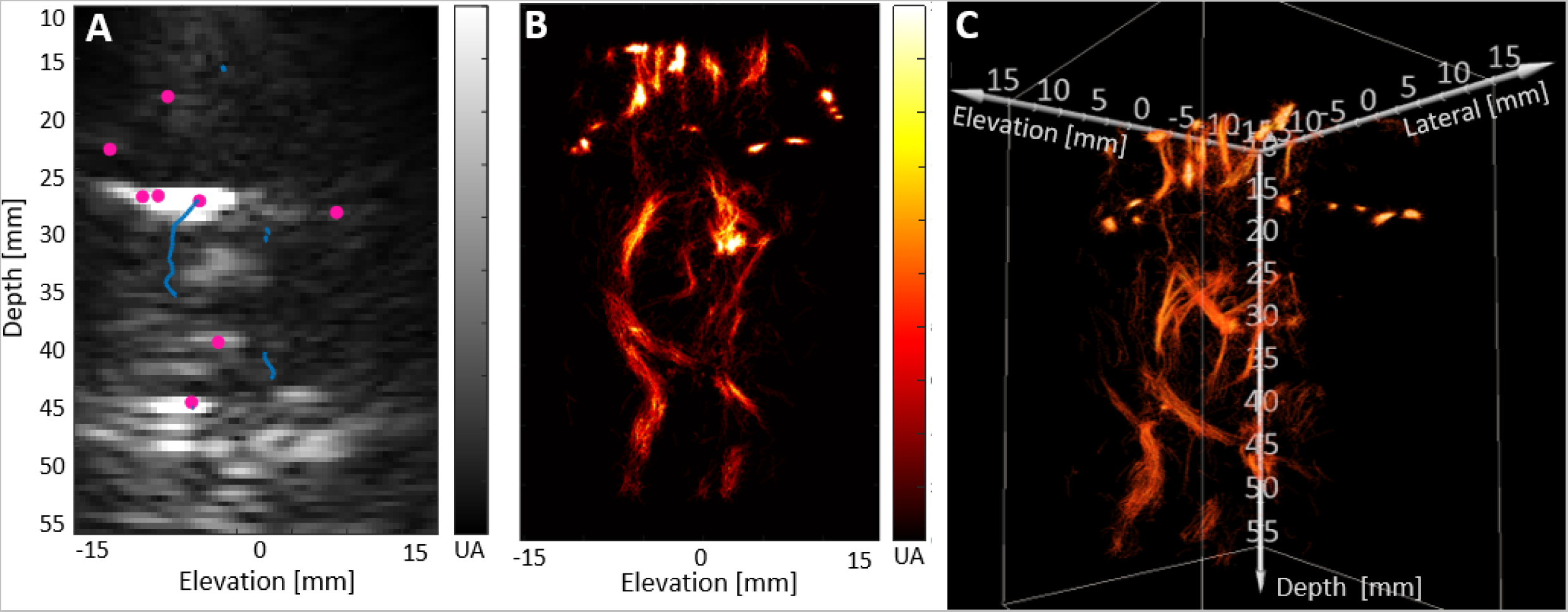
Examples of ULM processing in sheep M6 (1.75 ml/min). (A) Microbubbles (pink dots) and tracks (blue lines) on a 5 mm thick slice of filtered ultrasound imaging. Same sagittal view as in B., (B) Lateral projection of ULM density map (sagittal view), (C) Volumetric rendering of the same ULM density map with Amira software. The obtained ULM density map was then manually registered on the MRA volume using Amira software (Version 2019.4, Thermo Fisher Scientific, USA). The comparison of the vessels visible in ULM and in MRA was done by making a 5 mm thick slice with the help of a radiologist (S.B.)^58–60^.

#### b. ZTE MRI analysis

We manually synchronized ZTE MRI and MRA, allowing identification of the selected cranial zone through which ULM was performed. Several steps were realized to obtain skull thickness and attenuation defined by Guo et al.^47^ (Supplementary Fig.1). All these treatments were based on different assumptions. Firstly, we hypothesized a linear regression between logarithmic ZTE and CT-scan specifically for the skull bone^47^. Secondly, we exploited the CT-scan of one sheep to establish the linear regression between the two imaging modalities. This approach assumed that the imaging conditions were identical for all animal. As all acquisitions were performed in the same thermoregulated room, brain tissue and air were assumed constant on ZTE imaging for all animal,. Then, normalization was performed so that tissue was set to 2 and air to 1.

Statistical analysis of the skull attenuation was done using One-way ANOVA with GraphPad Prism (Version 9). Brown-Forsythe and Welsh ANOVA tests were performed assuming that standard deviations were not equal. The multiple mean comparisons were performed with Games-Howell test. Here, skull attenuation failed to pass normality tests but the high number of values in attenuation distributions (n>50000) allowed us to use ANOVA, i.e. n=50 563 for sheep M4, n=59 705 for M5 and n=62 391 for M6 with n the number of values in attenuation distribution for each sheep.

#### c. Micro-CT analysis

We analyzed separately, the diploë (manual segmentation with interpolation) and the inner and outer cortices segmented using the shrink-wrap method from CtAn (version 1.17.7.2). Bone mineral density measurements (BMD) were obtained after calibration of X-rays attenuation coefficients from bone based on the phantom calibration attenuation coefficients and were measured separately for diploë and both cortices.

Before morphological analysis, the bone structure was thresholded using the Otsu method (3d) for all datasets followed by opening morphological operations to remove noise (3d, radius=3). For the diploë, the following bone micro-architecture parameters were measured: total thickness (TT, mm), porosity which corresponds to Pore Volume over Total Volume (PoV/TV, %), trabecular thickness (Tb.Th, mm), trabecular separation (Tb.Sp, mm), trabecular number (Tb.N, mm^-1^), trabecular bone pattern factor (Tb.pfPf, mm^-1^), fractal dimension (FD), and the degree of anisotropy (DA). For the cortical layers, TT and porosity were measured. The Volume ratio in % of the different parts of the skull were computed from the mask. More details of the measurement methods are previously described in Chappard et al.^48^.

## Results

### A. ULM vs. MRI

Following ULM analysis, we were able to localize 1.3 + 0.5 10^4^ microbubbles per block of 400 volumes, in sheep M6 (injections of 1.5 ml/min). Tracking then yielded 35 ± 27 individual tracks per blocks, with a strong variability between animals and between blocks. From the resulting tracks, 3D density maps, i.e. count of microbubbles per voxel, or average velocity maps, i.e. average microbubbles speed per voxel, could be obtained (Fig. 2). ULM saturation curve, which represents the fraction of voxel occupied in the density map by at least one microbubble, increased naturally with the injected volume of microbubbles (Supplementary Fig. 3 C).

After manual co-registration, 3D ULM could be superposed on the MRA (Fig. 3). The ULM field-of-view remains limited to a volume of 3 × 3 × 6 cm^3^, which is larger than the probe size (Fig. 3 B). Within this limited volume, several blood vessels could be observed both on MRA and ULM. Some smaller vessels were uniquely detectable by ULM (Fig. 3 D-G). The precision of ULM allowed some vessels identification such as the internal cerebral vein. (Fig. 3 C-H).

**Figure 3.**
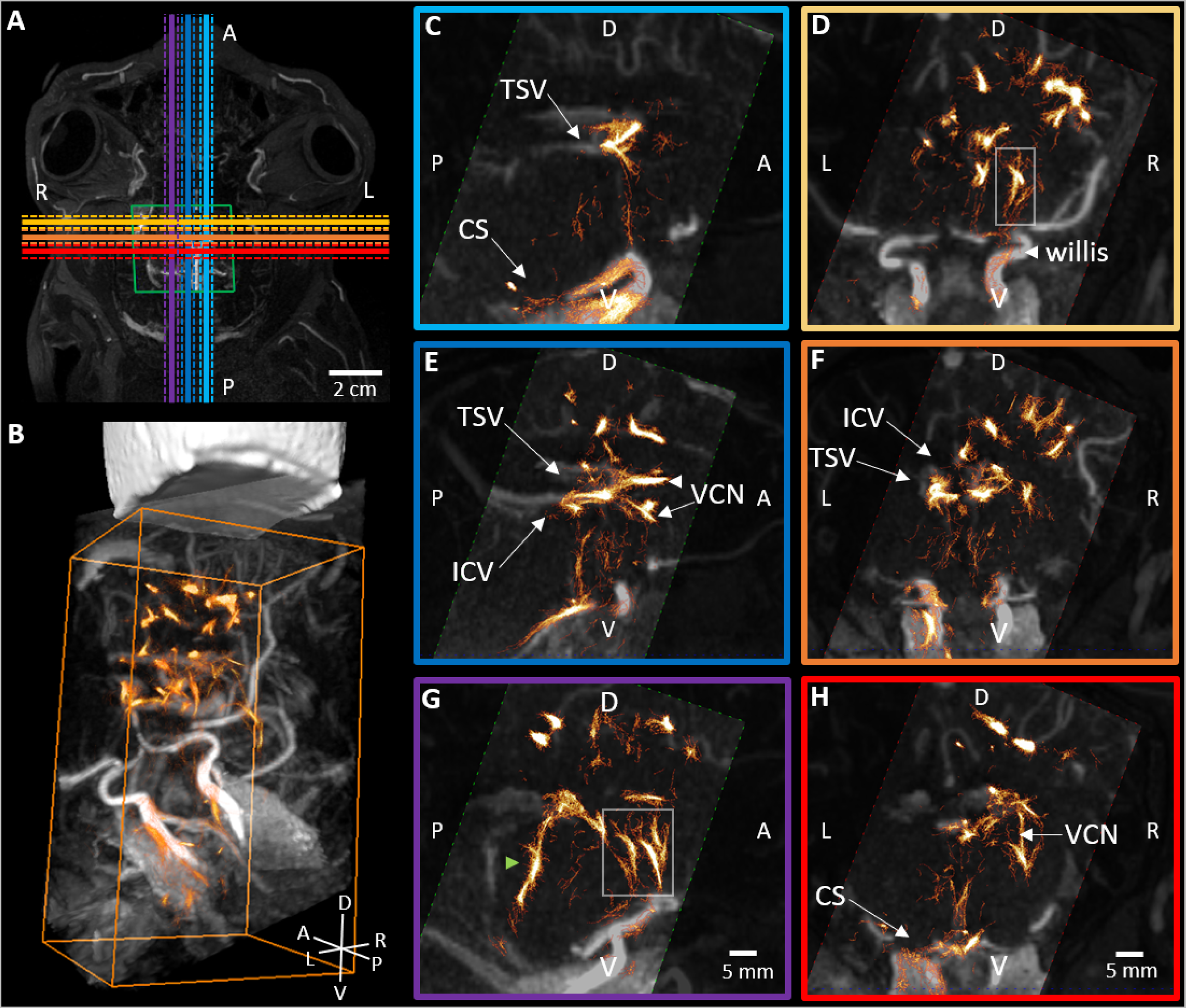
Superposition of ULM and MRA in M5. (A) Transversal slice of MRA with probe position (green box) and positions of different coronal or sagittal slices (purple to blue and yellow to red). The solid lines represent the middle of the slice and the dotted lines represent the 5mm thickness. (B) MRA and ULM in a volumetric view. The probe (in white) is placed according to the green box in A. (C, E, G) 5 mm thick sagittal slices. (D, F, H) Coronal slices. The image border color indicates the slice position in A. Green arrow indicates a shift between MRA and ULM. (G) White squares show vessels which are on ULM and not in MRA (D, G). Same scale for C to H. ICV: internal cerebral vein, TSV: thalamostriate vein, VCN: vein of the caudate nucleus, CS: cavernous sinus.

The circle of Willis and the principal veins were recognizable on the sagittal sections both on ULM and MRA (Fig. 4, Supplementary Fig. 2), while the Power Doppler only allowed the observation of larger vessels. (Fig. 4 D). Nevertheless, discrepancies were visible between the ULM and the MRA: in particular, a systematic offset was present between vessels (Fig. 4 A, Supplementary Fig. 2). Finally, some segment of vessels were not fully visible on the ULM or even did not appear at all. Due to a limited sensitive area, only a fraction of the Circle of Willis of the sheep was observable.

**Figure 4.**
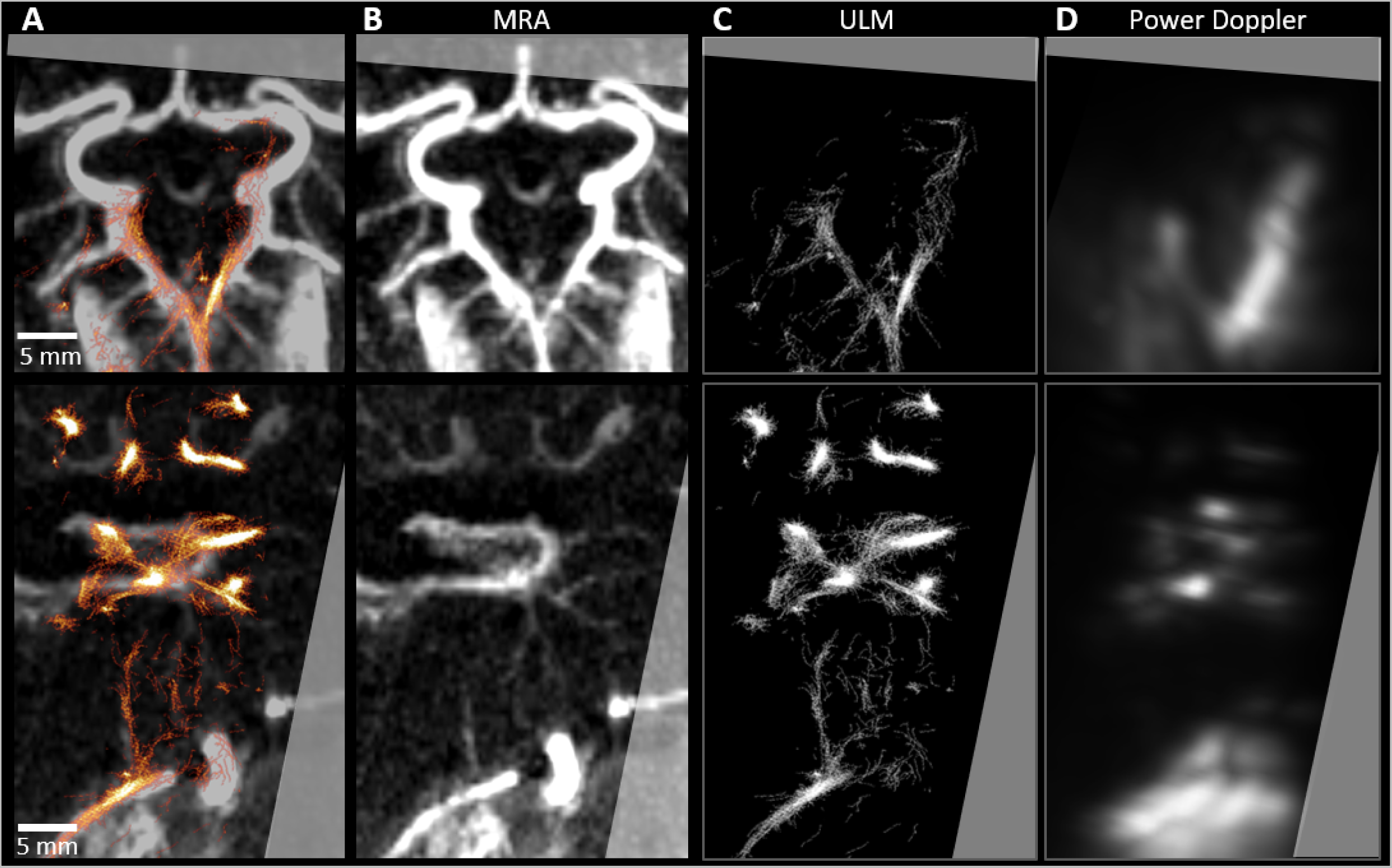
Comparison of ULM, MRA and Power Doppler. (A) Circle of Willis with MRA (gray colormap) and ULM (red colormap). (B) MRA only, (C) ULM, (D) Power Doppler. Thick slice of 5 mm. Coronal view on top, sagittal view on the bottom.

### B. ULM analysis

In this study, 3D ULM displayed a limited number of tracks, reducing the sensitivity to the microcirculation (Fig. 4). By making 1 mm sections of different ULM vessels in the ULM density maps (Fig. 5 A, vessels 1 to 3), the size of the observed vessels varied between 0.5 mm and 2 mm in diameter (Fig. 5 B). Only section 1 showed a Poiseuille profile, while the other vessels were aliased, such as the third one. Velocities measured within these vessels were between 20 and 200 mm/s, with most vessels having velocities around 60 mm/s. (Supplementary Fig. 3 A).

**Figure 5.**
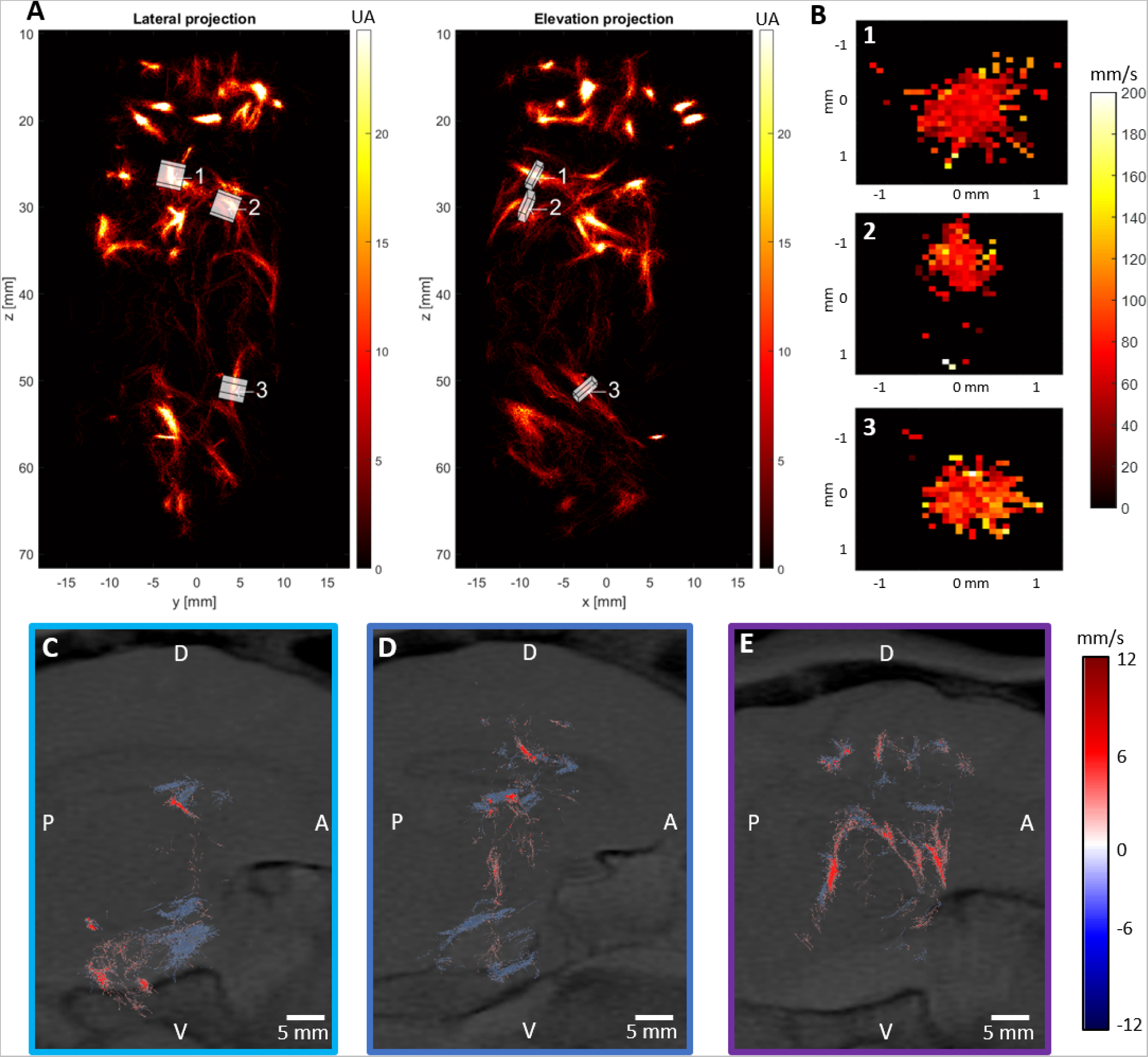
ULM characterization. (A) Positions of vessels section in ULM volume. (B) 1 mm thick slices vessels. (C, D, E) 5 mm thick sagittal slices of oriented velocity maps, same positions as in Fig. 3 C-G. ULM is overlapped with T2 MRI. Colormap shows the directivity of microbubbles speed: red for flow toward the probe, blue flow for flow away from the probe.

Based on the 3-dimensional displacement of the individual microbubble, it was possible to determine blood flow orientation in vessels and highlight different type of arteries or veins. For instance, blood was moving away from the probe in the case of the central veins (Fig. 4 C-D). On the other hand, the flow was going towards the probe in the case of the arteries originating from the Circle of Willis (Fig. 5 E). On few large vessels with high velocity flows, aliasing was visible. (Fig. 5 C-E)

### C. Skull analysis

The characterization of the skull properties with micro-CT was performed on the 3 sheep. Probe positioning varied between animals. For instance, M5 was imaged from a more caudal area than other sheep (Fig. 6 A-C). However, the skull sample for micro-CT scanning was systematically taken underneath the probe placement. The same portion of the skull in the axis of the probe was studied with ZTE MRI imaging.

**Figure 6.**
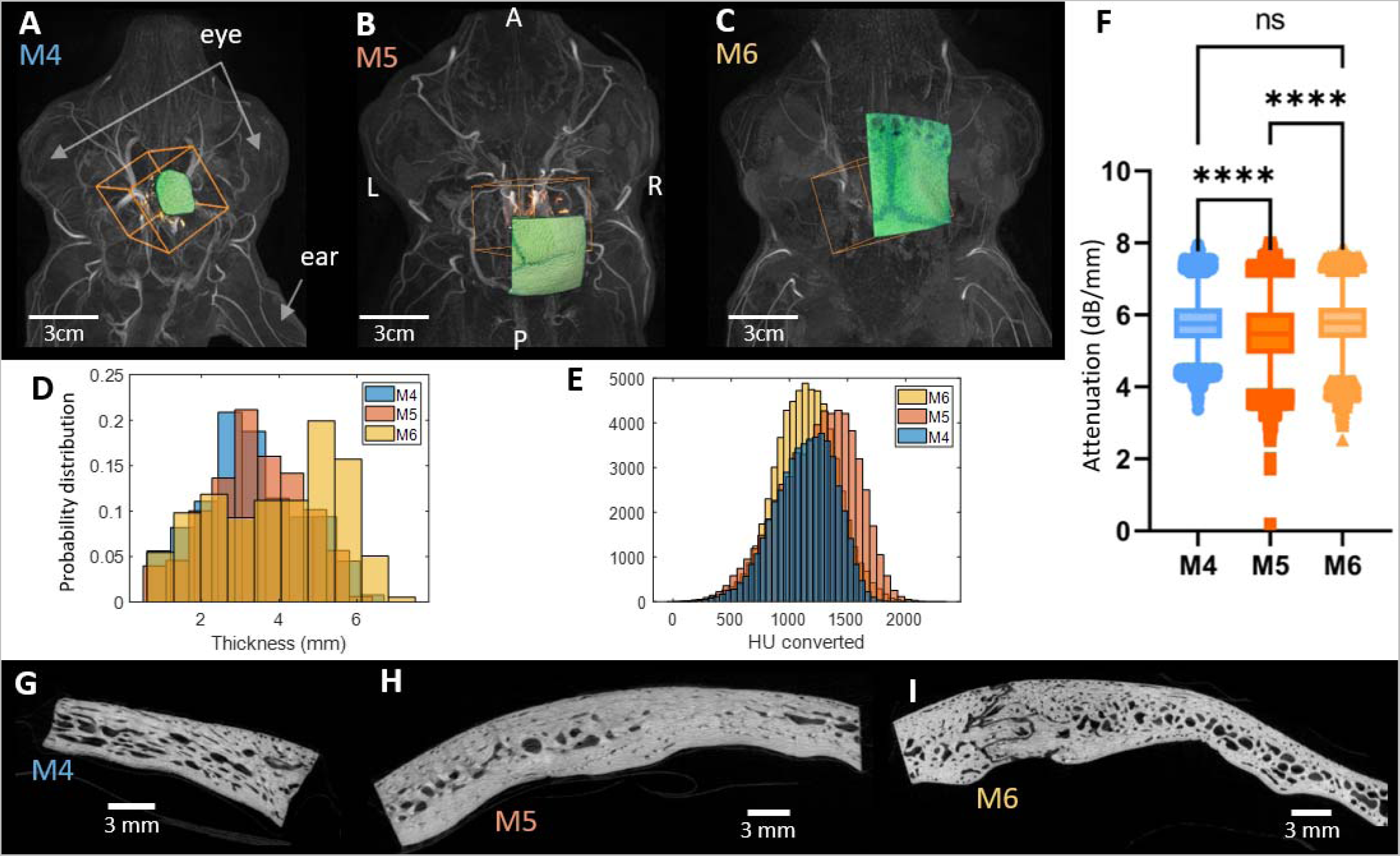
Skull bone analyses with micro-CT. (A) MRA co-registered with ULM (orange box) and with extracted skull sample imaged by micro CT (green) for M 4, (B) M5, (C) M6. (D) Thickness probability distribution measured on ZTE MRI. (E) Distribution of Hounsfield unit, HU, converted from ZTE MRI in the segmented skull. (F) Boxplot of attenuation computed from HU converted from ZTE MRI. (G) Skull bone slice from micro-CT volume of skull sample from M4, (H) M5, (I)M6. ****: p-value<0.0001, n.s.: non significative

The thicknesses assessed in ZTE MRI appeared to be more important for M6, with a mean thickness of 6 mm and an important variability. M5 and M4 thickness were more uniform with an average of 3 mm (Fig. 6 D).

The level of absorption measured in HU converted from the ZTE MRI shows different values for the 3 sheep, with a larger X-rays attenuation in M5 compared to M4 and M6 (Fig. 6 E). The conversion to acoustic attenuation at 1.5 MHz shows equivalent attenuations for the M4 and M6 sheep and a lower attenuation for the M5 sheep (Fig. 6 F).

The internal bones structure seemed different for M6 and M4 compared to M5 (Table 2). Indeed, the diploë volume ratio was much higher in the M4 and M6 sheep than in the M5 sheep. The inner and outer cortices of the sheep M5 correspond at least 1.5 time the cortices thicknesses of M4 and M6. The greater internal cortical bone thickness confirmed this in M5 sheep. Indeed, in some areas, the M5 sheep had no diploë at all. It was visible on the images (Fig. 6 G-H-I) where the porosity level of the diploë was more obvious for M4 and M6 then for M5. That phenomenon was also visible in BMD measurements representing the combination of bone material and mineral in the skull (Table 2). Indeed, a greater BMD of the cortical part was found in the M5 compared to M4. The M6 seemed to have the least dense cortical part.

**Table 2.** Micro CT result for sheep M4, M5, and M6. Inner: Inner Cortical bone, Outer: outer cortical bone, (standard value)

|  | Volume ratio (%) |  |  | Thickness (mm) |  |  | Porosity (%) |  |  | BMD gHA/cm<sup>3</sup> |  |  |
| --- | --- | --- | --- | --- | --- | --- | --- | --- | --- | --- | --- | --- |
|  | Inner | Outer | Diploë | Inner (STD) | Outer (STD) | Total (STD) | Inner | Outer | Diploë | Inner (STD) | Outer (STD) | Diploë (STD) |
| M4 | 19 | 33 | 48 | 0.6 (0.2) | 1.1 (0.3) | 4.2 (1.1) | 6 | 7 | 35 | 1.6 (0.5) | 1.6 (0.5) | 0.6 (0.5) |
| M5 | 36 | 36 | 28 | 1.8 (0.8) | 1.5 (0.5) | 5.3 (1.0) | 2 | 1 | 31 | 1.9 (0.3) | 1.9 (0.3) | 0.7 (0.5) |
| M6 | 14 | 29 | 57 | 0.7 (0.3) | 1.2 (0.3) | 5.7 (1.7) | 2 | 4 | 39 | 1.1 (0.1) | 1.1 (0.1) | 0.6 (0.2) |

## Discussion

The objective of this study was to demonstrate the potential of 3D ULM to image cerebral blood vessels through a skull comparable to human temporal bone. To achieve this goal, we used a low frequency multiplexed matrix probe connected to an ultrafast programmable ultrasound scanner able of achieving more than 200 volumes per second to detect and track in 3 dimensions the displacement of injected microbubbles. The sheep model was selected for the thickness of its parietal bone close to the human one. ULM were compared to MRA and skull bone characterization was done by ZTE MRI, CT and micro-CT imaging.

Cerebral blood vessels were observed beyond the sheep skull with 3D ULM and their location was overprinted on those observed in MRA. Nevertheless, the field-of-view of 3D ULM was restricted to a zone of about 3 × 3 × 6 cm^3^, corresponding to a divergent beam. This limitation could come from the skull shape, which is not very flat inducing ultrasound refraction. The microcirculation in the sheep brain observed by 3D ULM was thus much less apparent than in small animal models^43^. This is linked to the lower number of trackable microbubbles, despite the high and variable quantity of microbubbles injected compared to typical contrast-enhanced ultrasound in humans, i.e. 1 to 2 ml single bolus injection in humans against 1.5 to 2ml/min in sheep^49^. This could also be explained by the attenuation of the skull, which prevents 3D ULM from observing many microbubbles due to their low signal-to-noise ratio. For instance, M5 imposed less attenuation than M6 and, consequently, showed more microbubble tracks for the same volume of injections (Supplementary Fig. 3 B).

However, the absence of many microbubbles could also be explained by an anatomical filter at the basis of the sheep brain: contrarily to humans, sheep have a Rete Mirabile which is a capillary bed skull responsible for chemical and thermal exchanges of the blood before the irrigation of Wills circle^50,51^. The effect of such capillary structure on the microbubble is, to our knowledge, unknown but could be important. It should also be noted that there are no other large animal models available in which this phenomenon does not exist. The first three sheep were thus used to better define the quantity of microbubbles needed to obtain signal, such experimentation with the injection rate was necessary to adjust an adequate amount of microbubbles (Supplementary Fig. 3 C).

The slight deviation on the images between MRA and 3D ULM could have originated from aberrations within the skull, corresponding to a variability in speed of sound from the skull compared to that of brain tissue. Besides correcting shift between the MRA and the ULM, an aberration correction could allow an increase in microbubbles SNR and the elimination of vessels duplication, improving the ULM as a whole. Future studies should implement aberration correction described in several articles^52,53^.

The appropriateness of the sheep as a model for humans can also be discussed. The thicknesses of sheep’s skull bones is about few mm and close to a human temple^33,45^. The attenuation model on the skull of sheep inspired by the method carried out on humans show comparable attenuation, i.e. 5.8 dB/mm sheep and >6 dB/mm for humans at 1.5 MHz^47,54^.

Beyond these issues, this study suffers several limitations. As the complete experiment was performed on only three animals, the observed tendencies need to be reproduced. The first three animals allowed us to establish the experimental protocol, i.e. probe positioning and quantity of microbubbles to be injected. Due to the exploratory nature of this study, the experimental conditions varied between animals, including the attenuation of the skull, the positioning of the probe, the exact injected volume or certain acoustic conditions (Supplementary Fig. 4).

Based on a micro-CT study, in human parietal bone the inner cortical layer was about 0.5 mm thick and the outer one about 1 mm thick which was lower compared to values found in the present study for sheep^55^. Our results of diploe proportion in sheep between 0.28 to 0.57 are of the same order as the median value diploë proportion about 0.47 in humans^55^. Moreover, the inner and outer cortices present higher BMD and lower porosity in sheep than in humans^56^ (Table 2). The comparison at the microarchitecture level with the human parietal bone shows that the diploë of the sheep has a lower porosity and a higher BMD compared to humans^56^ (Table 2). The trabecular number (Tb.N) is similar, i.e. between 1.5 and 1.8 mm for sheep and about 1.69 mm-^1^ for humans^56^. Nevertheless, the architecture of the pores seems different between the sheep and the human, with larger Tb.Th for the sheep and lower Tb.Sp. (Supplementary Table 1). This difference in architecture is transcribed by very different Tb.Pf values between sheep and human, and a tendency of lower fractal dimension value in sheep than in humans. The negative Tb.Pf signifies a high connectivity with many concave surfaces and the FD is related to the surface complexity. The attenuation is lower in sheep compared to human and this can be explained by the fact that sheep were young subjects with not totally formed diploë, such as in M5. In summary, we can consider that the attenuation behavior of the sheep M4 and M6 will be more representative of the human condition. In addition, the temporal bone in the supra-aural zone in humans corresponds to the thinnest part of the skull and consequently should offer more favorable conditions for ultrasound propagation^57^.

To conclude, this exploratory study showed that the penetration-resolution trade-off could be alleviated by 3D ULM in large animal models, allowing the observation of cerebral blood vessels smaller than the wavelength. With the addition of aberration correction, further gain in resolution could be reached. Moreover, the human temporal bone being flatter than the sheep skull, and with less diploë, 3D ULM might be more favorable in homo sapiens than in ovis aries. However, imaging volume would still be restricted unless several imaging windows are used concurrently.

Although many improvements are still necessary, these results open the way for human studies with transcranial 3D ULM. The light ultrasound system designed in this study with a multiplexed ultrafast scanner allows to envision a wider use for cerebral imaging in the future in conjunction with large imaging system often overbooked (MRI, CT-scan). Not only does ULM achieve higher resolution, but the 3D nature of this approach permits the acquisition of large cerebral angiography easy to use without prior plane selection. Such angiography could, one day, help the triage of millions of stroke patients and improve the speed of care which is an important factor in the treatment success.

---

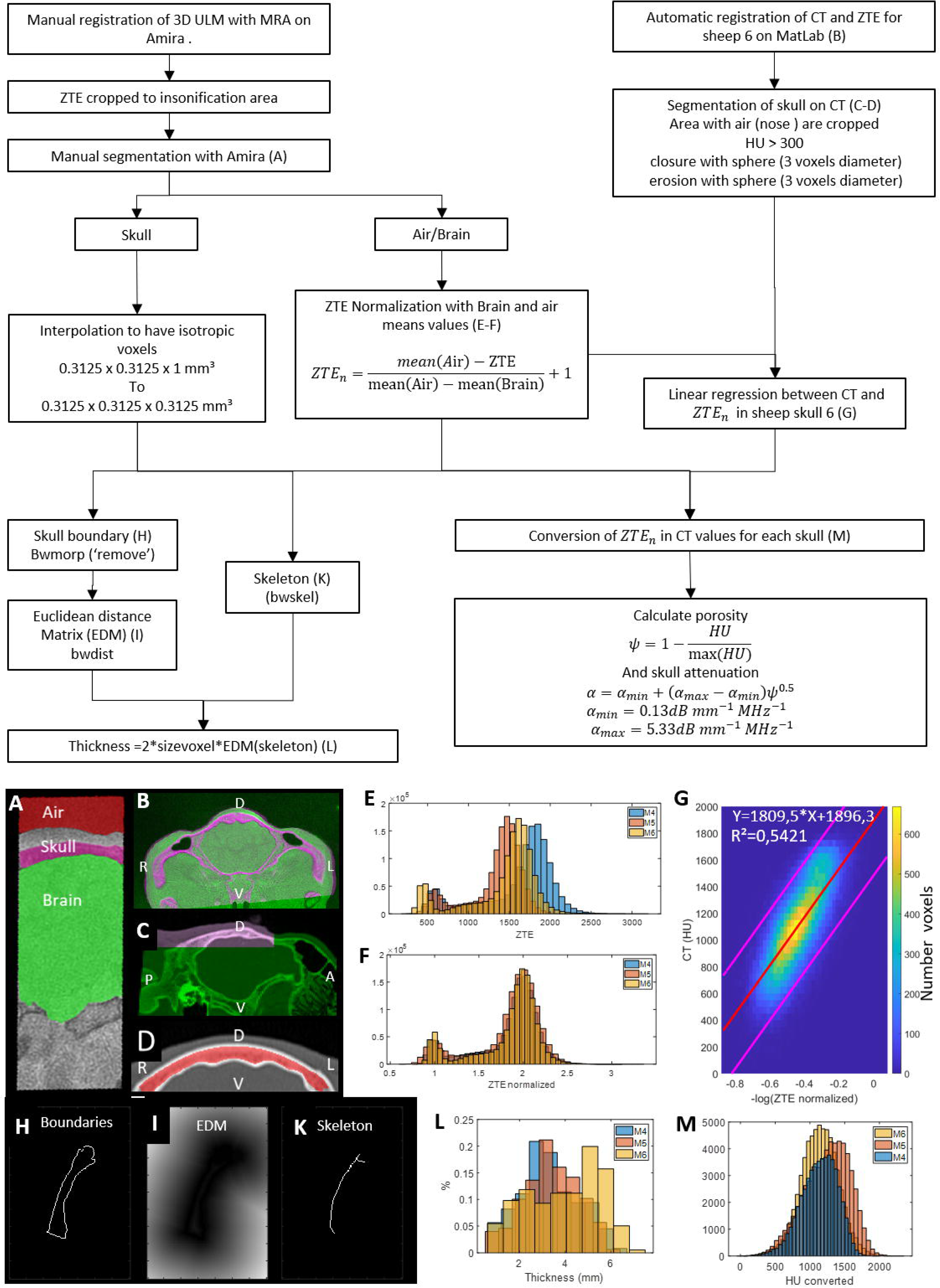

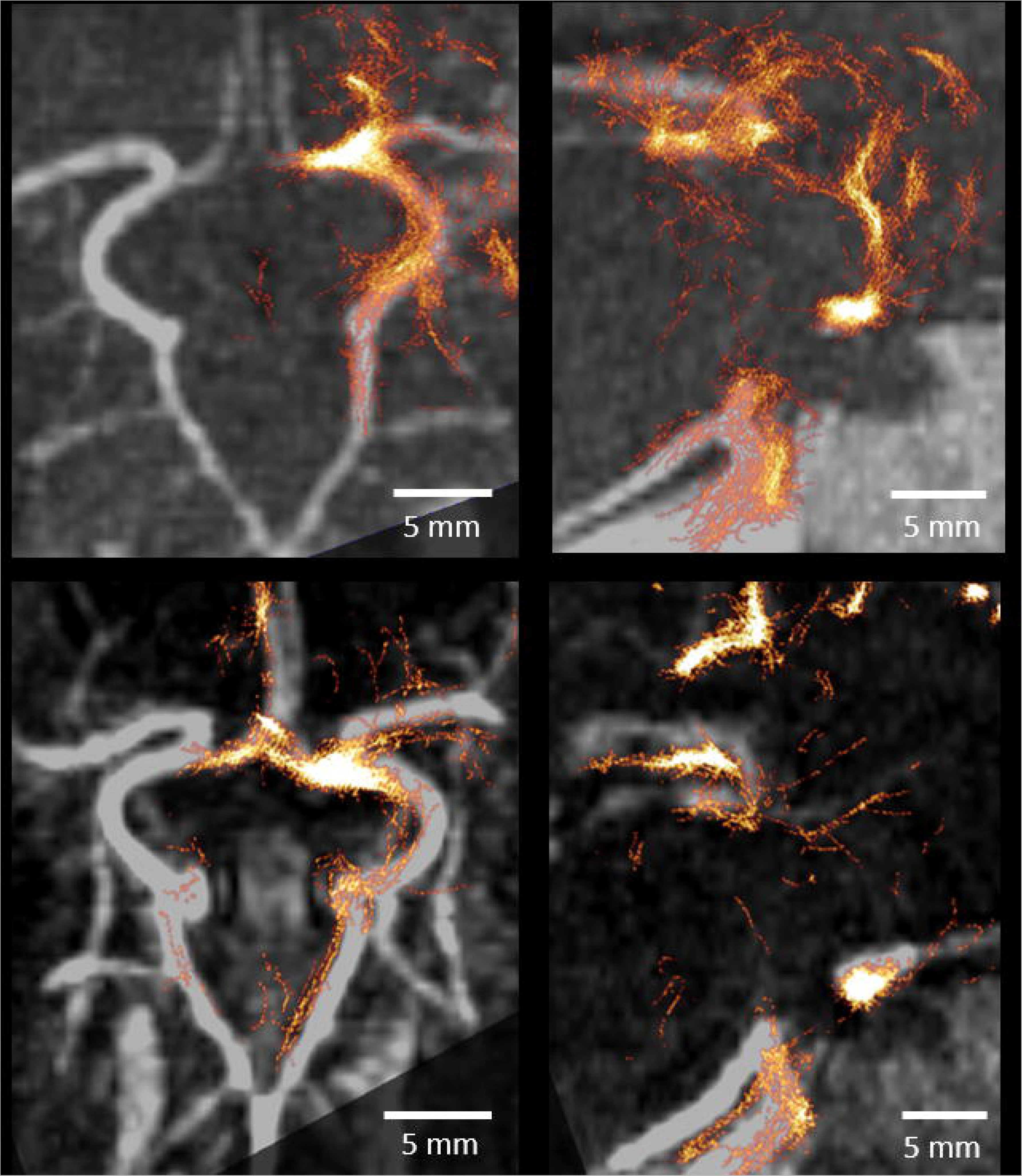

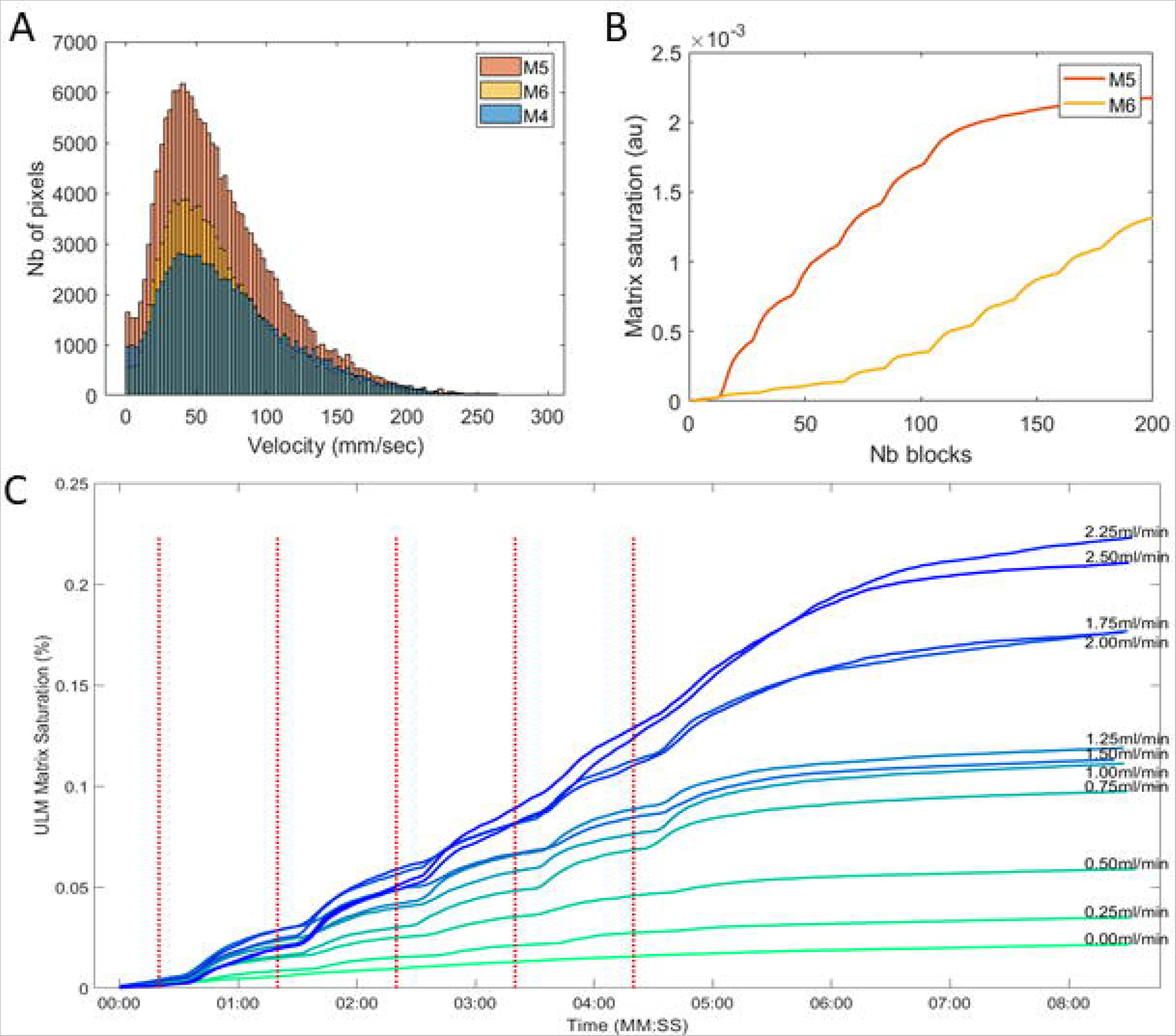

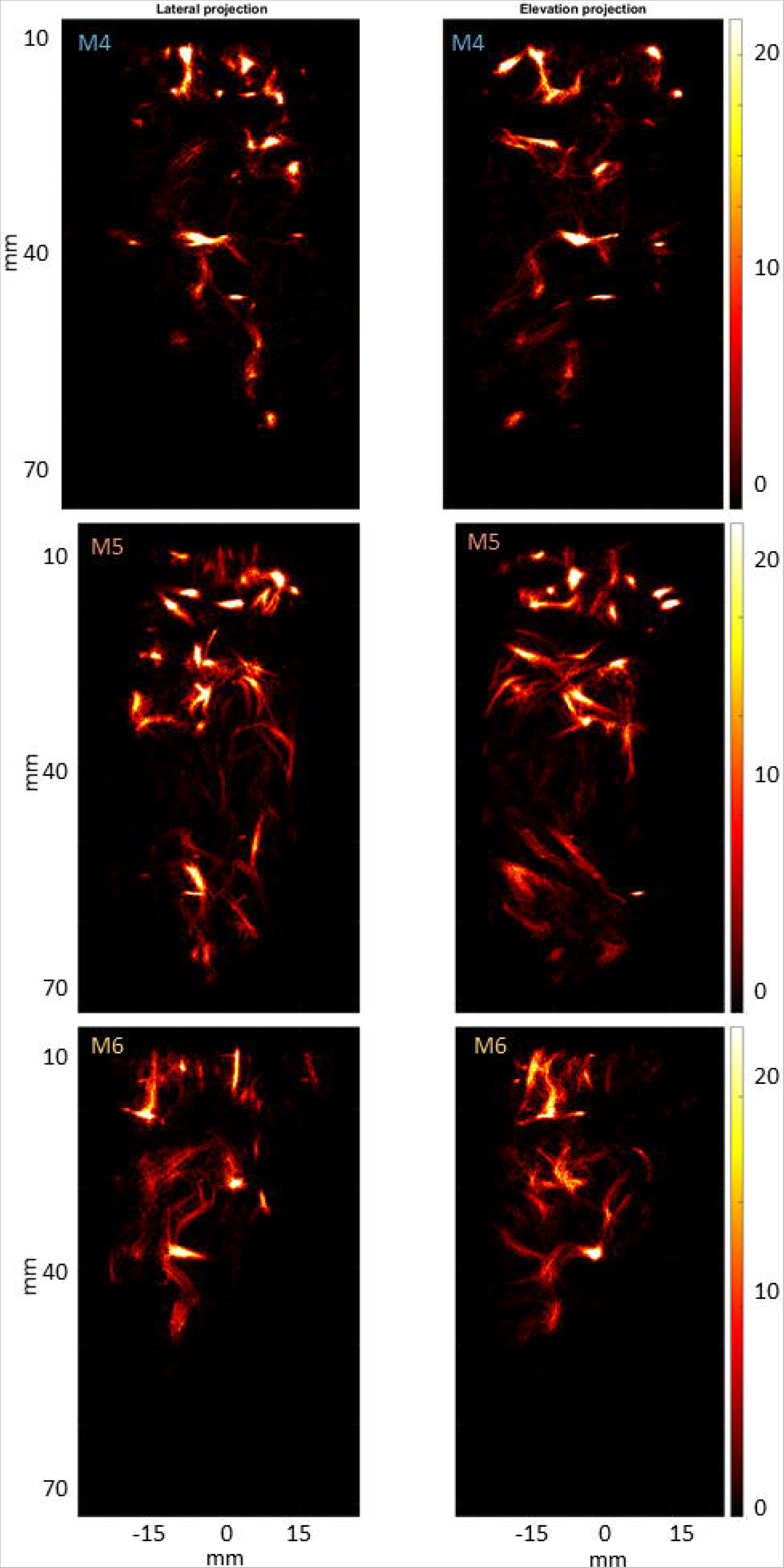

