## Supplementary table 1 and supplementary figures 1 to 4 for "3D Transcranial ultrasound localization microscopy reveals major arteries in the sheep brain"

### Supplementary materials

| Diploë | Tb.Th (mm)  (STD) | Tb.Sp (mm)  (STD) | Tb. N ($\boldsymbol{m}\boldsymbol{m}^{-\boldsymbol{1}}$) | Tb.Pf ($\boldsymbol{m}\boldsymbol{m}^{-\boldsymbol{1}}$) | DA  (STD) | FD |
| --- | --- | --- | --- | --- | --- | --- |
| M4 | 0,40 (0.14) | 0,44 (0.29) | 1.6 | -2.69 | 3.2 (0.7) | 2.51 |
| M5 | 0,37 (0.16) | 0.42 (0.24) | 1.8 | -3.62 | 2.3 (0.6) | 2.44 |
| M6 | 0,40 (0.14) | 0.48 (0.22) | 1.5 | -2.17 | 2.2 (0.5) | 2.55 |

**Table S1. Diploë parameters computed from micro CT** (standard value).

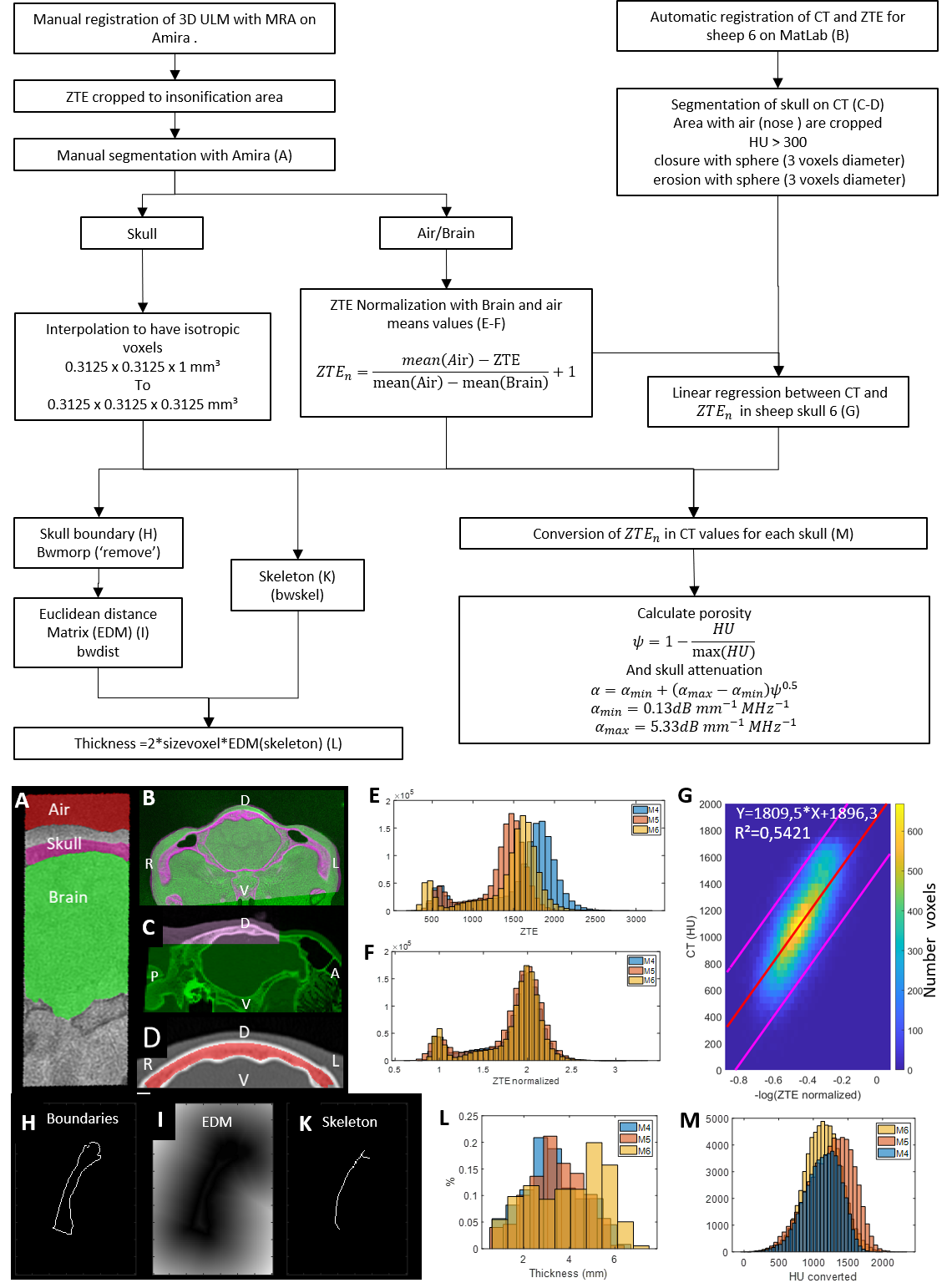

**Figure S.1 Schematic of steps for ZTE analysis** (A) Manual segmentation of ZTE on Amira (red: air, pink:skull, green:brain), (B) registration of ZTE and CT (same color encoding), (C) noise removing and skull/brain segmentation in another slice (same color encoding), (D) skull segmentation on CT in the same slice (red: skull segmented). (E) ZTE distribution before normalization for the three sheep, and (F)after normalization. (G) Linear regression between CT and -log ZTE. (H) Boundary of the skull. (I) Euclidian distance matrix, (K) skeleton of the skull boundaries according to Guo et al^47^. (L) Distribution of skull thickness for the three sheep. (M) Distribution of CT converted value from ZTE for the three sheep.

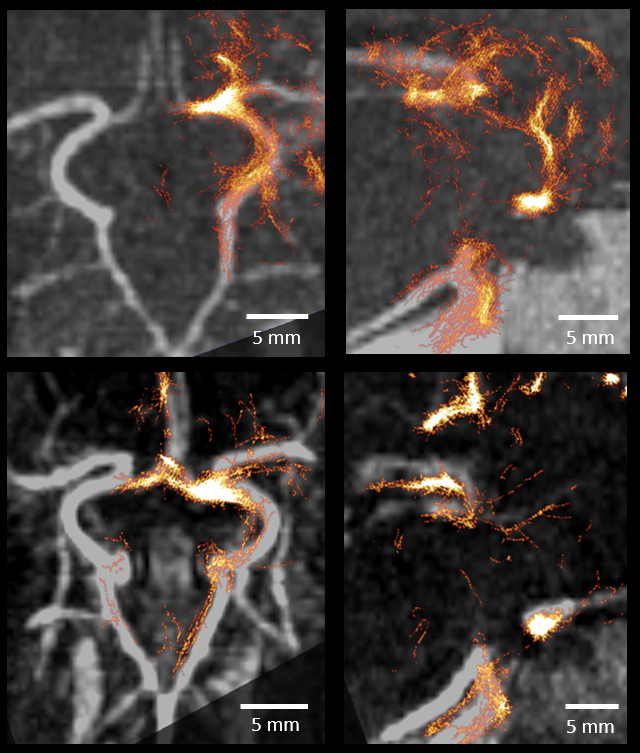

**Figure S2. Unilateral ULM overlapped on bilateral MRA, and unilateral ULM overlapped on bilateral CT-angiography in M6**. On the top: Willis circle in a 5 mm thick slice with MRA (gray colormap) and ULM (Orange colormap), On the bottom: same 5 mm thick slices with CT-angiography and ULM.

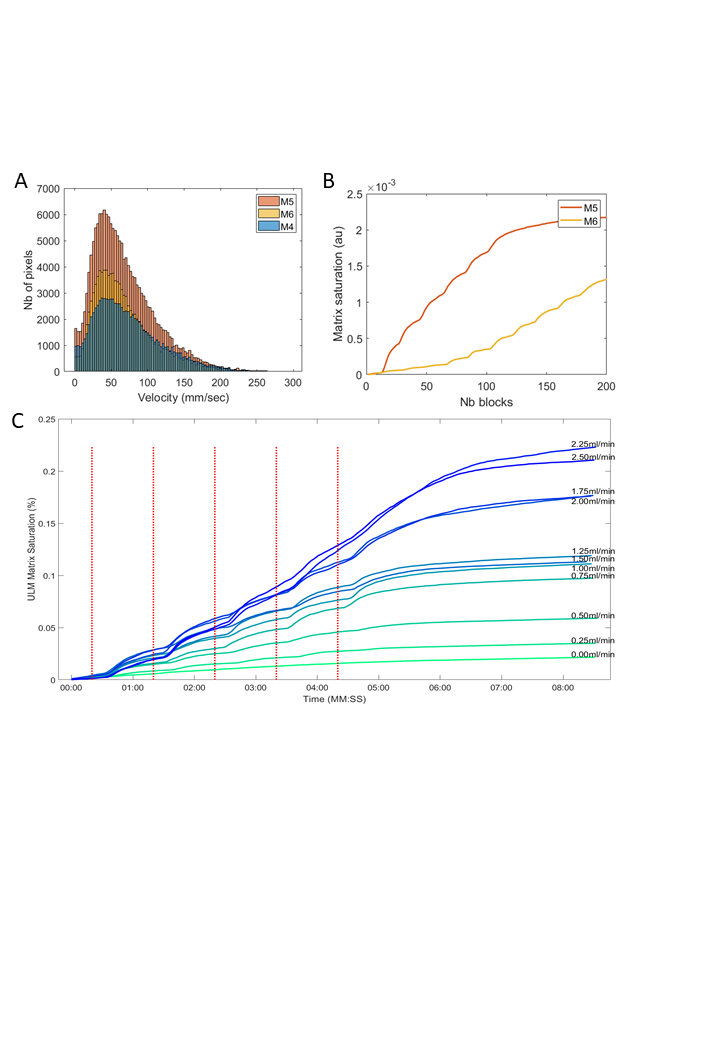

**Figure S3. Analyses of velocity, saturation and injection**. (A) Microbubble velocity distribution. (B) ULM saturation curves for M5 and M6. (D) ULM saturation for different kind of volume of injection on sheep M6.

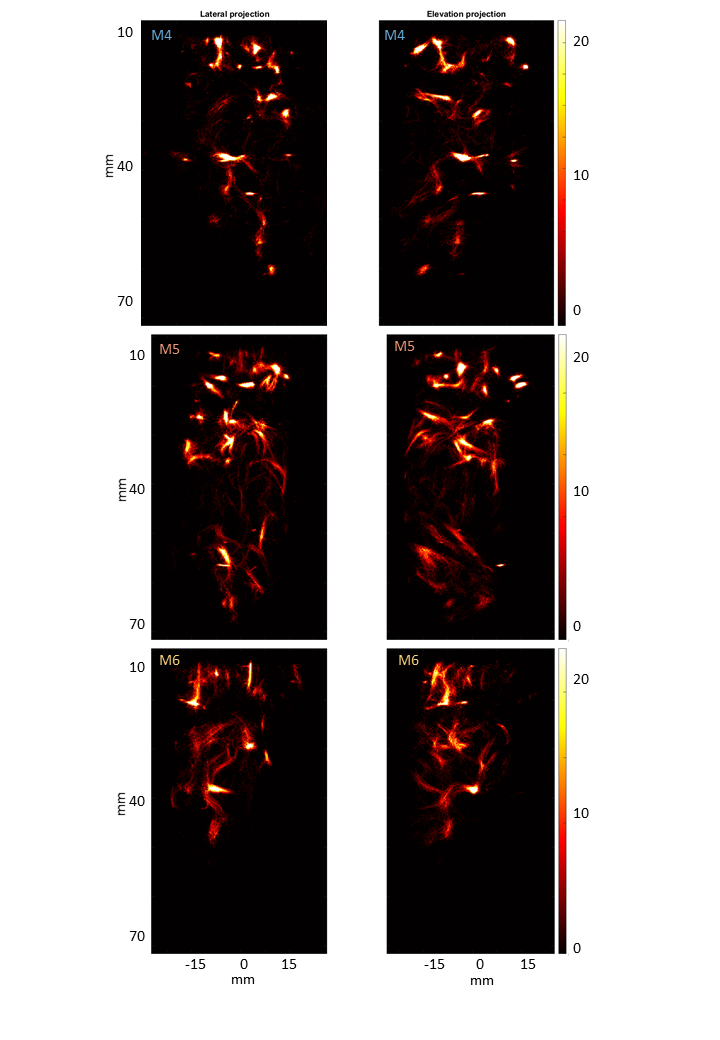

**Figure S4. ULM density maps**. From top to bottom: M4, M5, M6. Colormap are on arbitrary units.
